# Functional insights into the photoactive yellow protein family from homologs, multidomain proteins and inferred *pyp* operons

**DOI:** 10.1101/2025.06.04.657921

**Authors:** Rosalie L. Dohmen, Gunnar Hoogerwerf, Amber J. Dohmen, Madison L. Neal, Catalina Bradley, Clarice Huffman, Sarah M. Teeman, Saylor F. Hampton, Scout Powell, Zariah Gunn, Wouter D. Hoff

**Affiliations:** Department of Microbiology and Molecular Genetics, Oklahoma State University, Stillwater OK 74078, USA

## Abstract

Photoactive Yellow Protein (PYP) is a model system for functional protein dynamics and a prototype of the PAS domain superfamily. It is a bacterial photoreceptor that triggers a range of responses in different bacteria: phototaxis, biosynthesis of photo-protective pigments, and light regulation of biofilm formation. An important gap in knowledge on PYP is the signal transduction chain that guides the initial signal from the photoreceptor to various biological responses. Here we report an expanded set of 984 PYP homologs, providing information on sequence conservation and variation. We analyze this set of PYPs using two bioinformatics approaches to identify candidate proteins that are functionally related to PYP. First, we identified 153 multi-domain proteins containing PYP and analyzed the domain composition of these proteins. Specific preferences for N- or C-terminal placement of the PYP domain were observed. Second, we identified 113 predicted multi-gene operons containing the *pyp* gene. These two approaches yielded multiple candidates for proteins in the signal transduction chain associated with PYP, particularly histidine kinase (implying phosphorylation), methyl accepting chemotaxis protein (implying phototaxis), and GGDEF and EAL proteins (implying a role of c-di-GMP and biofilm formation). Some of these candidates were present only in multi-domain proteins and others only in *pyp* operons. Overexpression of the PYP domain from the MCP-fusion protein from *Nitrincola alkalilacustris* yielded a protein with an absorbance maximum of 447 nm and an overall photocycle rate of 0.5 seconds. Our results provide a clear basis for future experimental work on identifying signal transduction partners of PYP.

## Introduction

Evolutionary diversification causes protein sequences in nature to occur in protein families and often is accompanied by functional diversification. Pfam is a major database aimed at organizing the ever-increasing number of protein sequences derived from genome sequencing into such protein families (1). In addition, members of protein families, or domains in these proteins can be combined into myriad different combinations based on domain shuffling, leading to novel functions by functional coupling between such domains (2). The identification of such conserved domains has resulted in databases that allow proteins to be analyzed in terms of their domain composition, often resulting in insights regarding their possible biological function. Thus, diversification of protein families and domain shuffling are major drivers of evolutionary innovation at the protein level (3).

The proteins that belong to a protein family generally share a range of properties. They tend to conserve key aspects of their function and mechanism, including a conserved active site, a common three-dimensional structure, and a characteristic pattern of additional conserved residues. All members of such a protein family derive from a common ancestral protein that first evolved this combination of functional and structural properties. A classic example of such a protein family are the serine proteases (4). Here we study the photoactive yellow protein (PYP) family of photoreceptor proteins (5–12).

One important evolutionary process extends beyond this description. During the evolutionary diversification of a protein family, a shift in functionality can occur (neofunctionalization), in which the protein evolves to acquire a novel function, often resulting in loss of otherwise functionally critical conserved residues. As such neofunctionalization proceeds, a highly diverse set of proteins can emerge that still share a common three-dimensional structure and (often very weak) patterns of sequence conservation. However, these proteins no longer share a conserved active site and often are functionally highly diverse. Such a group of proteins is referred to as a protein superfamily. An important example is the Per-Arnt-Sim (PAS) domain superfamily (13–15). PYP is a member of the PAS domain superfamily and at the level of biophysics is the most deeply studied member of this superfamily.

The biochemical and biophysical properties of the founding member of the PYP family, isolated in 1985 (9, 10) from the halophilic and photosynthetic purple sulfur bacterium *Halorhodospira halophila*, have been studied extensively (5–8, 12, 16). The PYP from *H. halophila* (Hhal PYP) consists of 125 residues and exhibits an absorbance maximum (λ_max_) near 447 nm caused by its deprotonated *p*-coumaric acid (*p*CA) chromophore (17–19) covalently bound to Cys69 via a thioester bond (20, 21). The crystal structure of PYP revealed Tyr42 and Glu46 as key active site residues that interact with the *p*CA chromophore (22). Photoexcitation of PYP triggers a multi-step photocycle (10, 23) that is initiated by *p*CA photoisomerization (24, 25) to yield a red-shifted photocycle intermediate. This is followed by proton transfer from the protonated side chain of Glu46 to the deprotonated phenolic oxygen of the *p*CA (25), which triggers large protein conformational changes, including partial protein unfolding (26–32), resulting in a blue-shifted pB photocycle intermediate that decays to the initial dark stake with a lifetime (τ_pB_) of approximately 0.5 seconds. The biological function of Hhal PYP is to trigger negative phototaxis towards blue light (33), and the pB intermediate is the likely candidate for being the activated form of PYP that serves as the signaling state for initiating phototaxis signaling.

Subsequent work has yielded biochemical information on 10 additional PYP homologs: from *Chromatium salexigens* and *Rhodospirillum salexigens* (34), from *Rhodobacter sphaeroides* and *Rhodobacter capsulatus* (35–38), from *Rhodospirillum centenum* (39–41), from *Thermochromatium tepidum* (42, 43), a second PYP homolog from *Halorhodospira halophila* (44), from *Salinibacter ruber* (8, 45–47), from *Idiomarina loihiensis* (48), and from *Leptospira biflexa* (49). In parallel, bioinformatics analyses have yielded a rapidly growing number of proteins with amino acid sequences that identify them as PYP homologs, from 14 (8) to 134 (11) to most recently 789 PYP homologs (50).

Research on this growing set of PYP homologs clearly illustrates the phenomena of active site and mechanistic conservation, functional diversification, and domain shuffling. With respect to sequence conservation, published work yielded a set of 9 residues forming a conserved *p*CA binding pocket consisting of Ile31, Tyr42, Glu46, Phe62, Val66, Ala67, Pro68, Cys69 and Phe96 (using the residue numbering of the PYP from *Halorhodospira halophila*), with 15 additional residues showing a high level of conservation in the PYP family identified as Leu26, Phe28, Gly29, Asp34, Gly37, Asn43, Gly59, Phe63, Phe75, Gly77, Phe79, Gly86, Phe92, Val105 and Trp119 (8, 11, 50). A high level of conservation of the three-dimensional structure in the PYP family was demonstrated by the crystal structure of the PYP domain from *Rhodospirillum centenum* (51). All of these 11 PYPs exhibit a light-triggered photocycle resembling that of Hhal PYP. However, the spectral, biochemical, and kinetic properties of these PYP homologs varied considerably (8, 34–49). A striking example of this functional variation is that the final, slowest step in the PYP photocycle (from the blue-shifted intermediate back to the initial state) occurs in approximately 1 second in the PYP from *Halorhodospira halophila* (10, 23), but approximately 1 hour in the PYP from *Salinibacter ruber* (46).

Information on the biological function of PYP is limited, but has also revealed variability in the *in vivo* functional output of PYP triggered by blue light excitation. While experimental results show that in *H. halophila* PYP triggers negative phototaxis (33), in *Rhodospirillum centenum* PYP causes induction of genes encoding enzymes involved in photoprotective pigment biosynthesis (39), and in *Idiomarina loihiensis* it causes a blue-light induced reduction in biofilm formation (48) (Figure 1). Further insights were obtained from analyses of the genetic context of the *pyp* gene. The gene encoding Hhal PYP is part of an operon that also encodes tyrosine ammonia lyase and *p*-coumaryl Coenzyme A ligase, the two proteins needed to synthesize and activate the *p*CA chromophore (37, 52, 53). Meyer et al., 2012 (11) extended the bioinformatics analysis of genes flanking *pyp* genes to infer further insights into the biological function of PYP, which will be discussed in more detail below.

**Figure 1.**
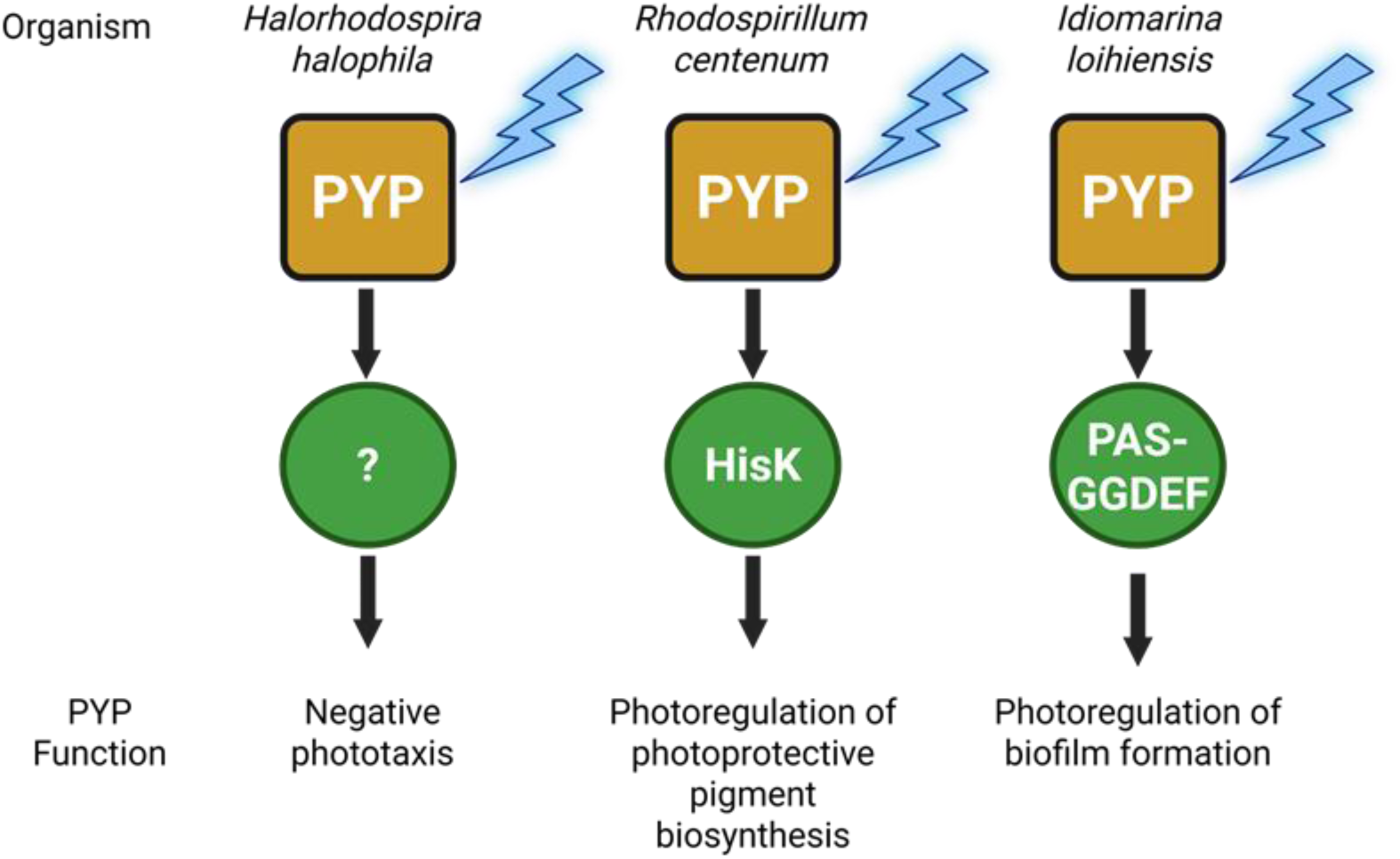
Summary of experimental data on the physiological functions initiated by PYP. Many aspects of the signal transduction chains involved remain unknown.

Regarding domain shuffling, all initially identified PYPs occurred as single-domain proteins approximately 125 residues in length. However, subsequent work revealed PYP domains in much longer proteins, leading to the identification of a quite small set of protein domains to which PYP can be fused in nature. The first of these multi-domain PYPs was found in *Rhodospirillum centenum*, where it is part of proteins also containing a light-sensing bacteriophytochrome (54) and a Histidine kinase that is regulated by photoexcitation of the PYP domain (39). Additional multi-domain PYP proteins were found in *Thermchromatium tepidum,* where the PYP domain is part of a protein also containing a bacteriophytochome and a GGDEF/EAL domain (42, 43). GGDEF/EAL proteins function in the biosynthesis and breakdown, respectively, of c-di-GMP, a signaling molecule that is closely associated with the regulation of biofilm formation (55). In addition, bioinformatics analysis revealed an MCP-PYP multidomain protein in *Methylomicrobium alcaliphilum* (11). Since methyl-accepting chemotaxis proteins (MCPs) regulate bacterial motility during chemotactic and phototactic responses (56) via an associated two-component regulatory system (TCRS) based on phospho- relay, a genetic linkage between an MCP and a PYP supports a possible role of this protein in phototaxis.

While a wealth of information is available on the biochemistry and biophysics of PYPs (5–8, 25, 28, 57), much less is known about its *in vivo* signaling. Here we report genes that are clear candidates for being functionally related to PYP in order to help identify the signal transduction chain of PYP and the biological functions triggered by members of the PYP family. We employed two strategies: analysis of the domain structures of multi-domain proteins containing PYP and predicted operons that contain the *pyp* gene. The first approach employs the phenomenon of domain shuffling, resulting in multi-domain PYPs in which PYP is part of a larger protein, and is based on the assumption that domains located within the same protein are functionally related. The second approach relies on the classic finding (58) that genes in bacteria often are arranged in operons, where functionally related genes are located immediately adjacent to each other and are co-transcribed into a single mRNA molecule. While their exact mechanistic implications remain actively debated (59–62), the occurrence of such operons is widely used to infer clusters of functionally related genes. The expanded PYP family studied here using these two approaches both validates and expands knowledge on the functioning of this family of bacterial photoreceptor proteins.

### Expanding the size and taxonomic distribution of the PYP family

In 2008, a total of 14 PYPs were identified across 12 distinct organisms, with the majority of these proteins being found within the phylum Pseudomonadota (8). At that time, it was noted that PYPs exhibit significant diversity, both in their functional properties and their amino acid sequences. A few years later, the number of identified PYPs had risen dramatically to 140, largely due to advancements in metagenomic analysis and whole-genome sequencing (11). These PYPs were derived from a broad array of bacterial species inhabiting various biological niches. In 2022, the number of identified PYP homologs expanded further, reaching a total of 789 sequences (50). In the present study, we report 984 PYP homologs. Technical details bioinformatics analysis procedures can be found in the supplement.

While photoreceptors and PAS domain proteins are found in the three domains of life (77), PYP has only been identified in bacteria. Most of the PYP homologs analyzed here are present in the Phylum Pseudomonadota (previously Proteobacteria) (67%), with representation distribution observed over all the classes of this phylum (Figure 2). Furthermore, PYP homologs were identified in the FCB (Fibrobacterota, Chlorobiota, Bacteroidota) superphylum (10.2%), with PYP present in the Bacteroidota, Gemmatimondota, and Rhodothermota. The Myxococcota contained 9.7% of the PYP homologs, while 5.9% were present in Spirochaetota. The remaining PYP homologs were present in the Acidobacteriota (2.4%), Terrabacteria group (2.3%), the PVC group (1%), and the Bdellovibrionota (1%). In the case of the PVC group (Planctomycetota, Verrumicrobiota, Chlamydiota), we found PYP homologs in the Planctomycetota and Verrumicrobiota. For the superphylum Terrabacteria we observed homologs in the phyla Candidatus Sericytochromatia, Candidatus Eremiobacterota, Armatimonadia and Actinomycetes. Finally, Thermodesulfobacteriota and Campylobacterota contained <1% of the PYP homologs.

**Figure 2.**
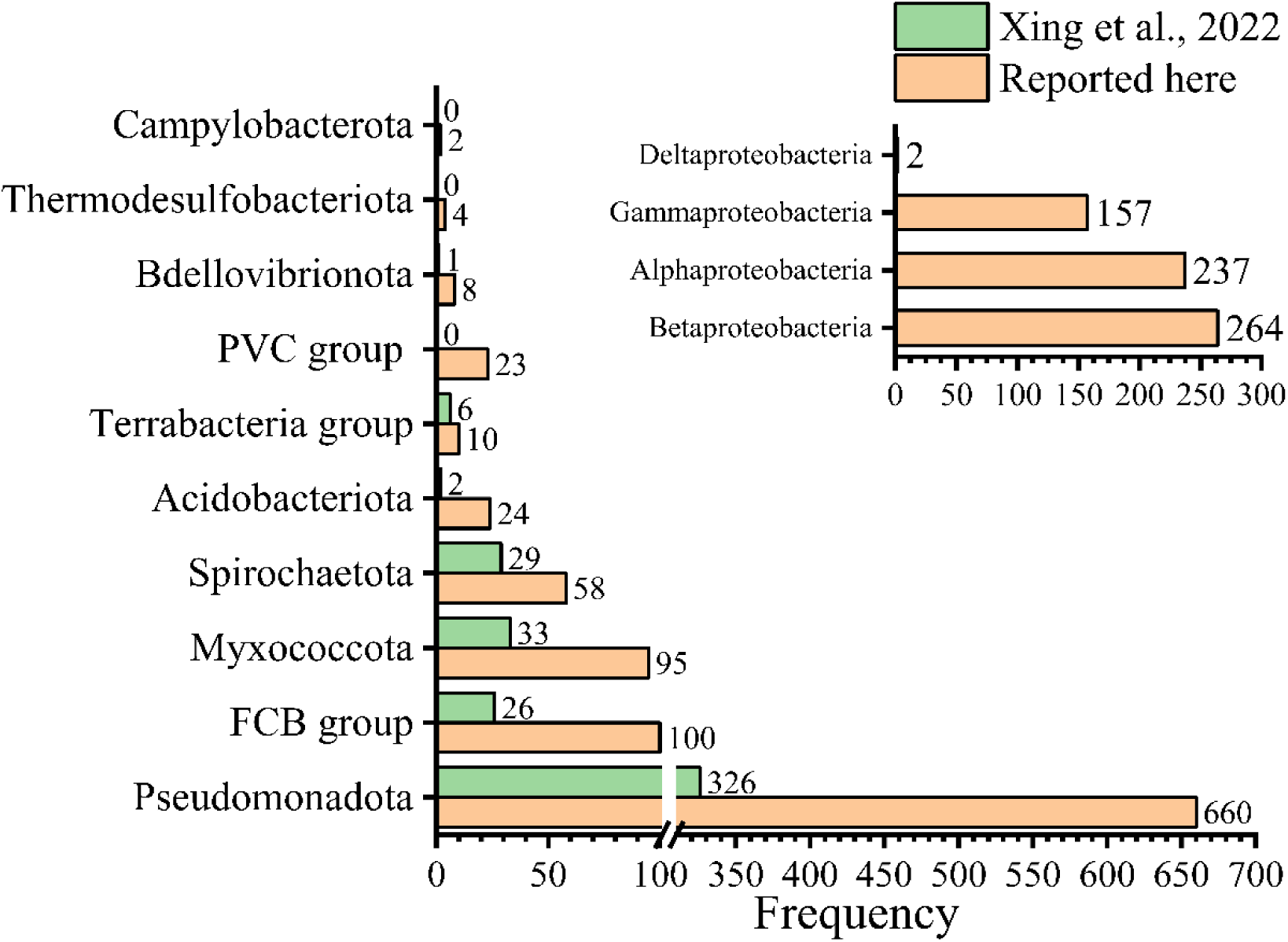
Distribution of taxonomic phyla that contain PYP homologs. The taxonomic data of 984 PYP homologs were extracted and analyzed. The Terrabacteria group, FCB group, and the PVC group are superphyla. The taxonomic distribution reported here (orange bars) is compared to that reported by Xing et al., 2022 (50) (green bars). The number of PYP homologs reported here for each taxon is indicated.

Taken together, these data confirm the taxonomic distribution of PYP across the bacterial domain as previously reported (50) and expand the presence of PYP homologs to three additional phyla.

### Sequence conservation and variation in PYP

The expanded set of PYP homologs reported here provides an opportunity to refine insights into the pattern of both sequence conservation and variation in the PYP family. To assess the degree of sequence divergence of the set of 984 Cys69-containing PYP homologs studied here, we computed the pairwise % sequence identity and % sequence similarity for each pair of PYPs (see (8)) and examined the distribution of the resulting similarity and identity levels (Figure 3). This analysis revealed a bimodal distribution of pairwise sequence identity values, with maxima near 39% and 58% sequence identity, and with % identity values dropping down to 27%. For sequence similarity a maximum value near 68% was seen, extending down to 44% similarity. These results demonstrate that the PYP family exhibits quite strong sequence diversity.

**Figure 3.**
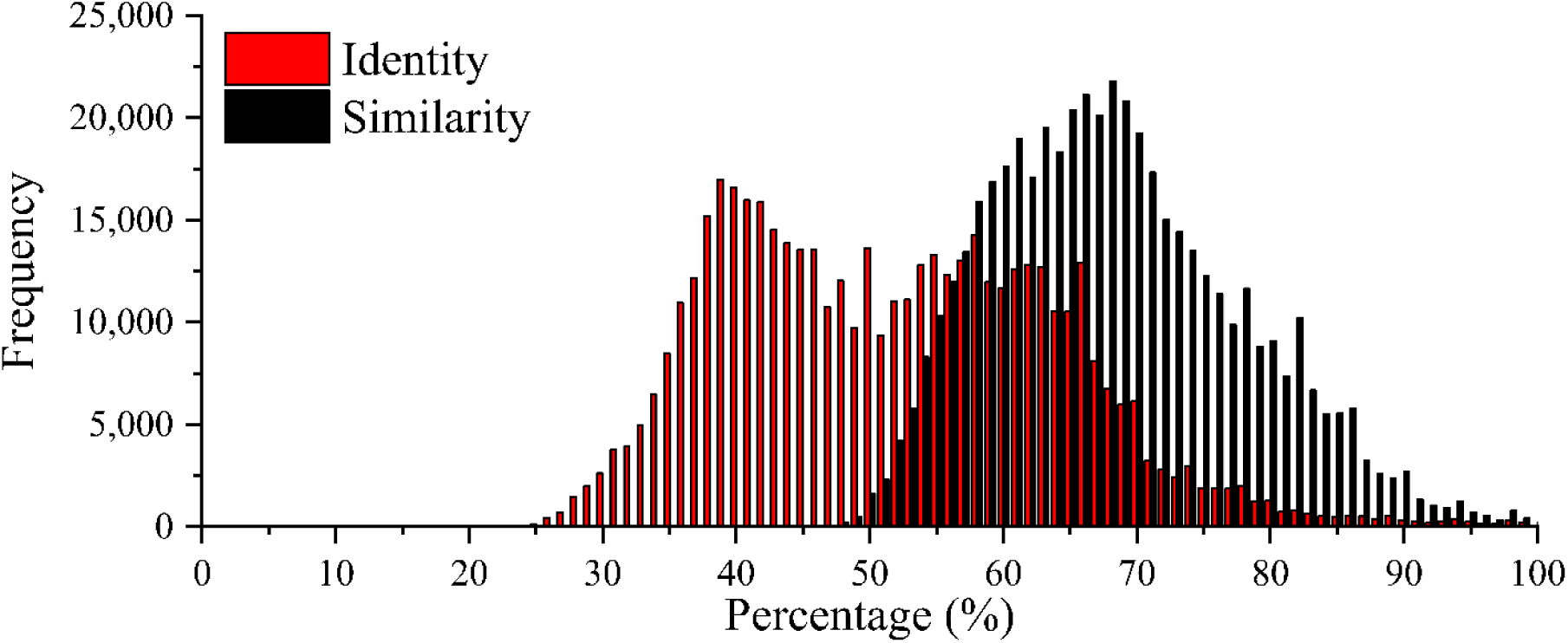
Distribution of the values of pairwise percentage identity and similarity of the 984 PYP homologs reported here. The percent identity is annotated in red, while the percent similarity is shown in black.

To gain insights into amino acid sequence conservation, a multiple sequence alignment (MSA) was computed and analyzed. In line with previous results (8, 11, 50) the set of 984 proteins studied here exhibited multiple strongly conserved residues and a high degree of conservation of the length of PYP near 125 residues, facilitating the process of obtaining a high-quality MSA. To gain insights from this MSA, we extracted the following information (Figure 4), using residue numbering based on Hhal PYP.

**Figure 4.**
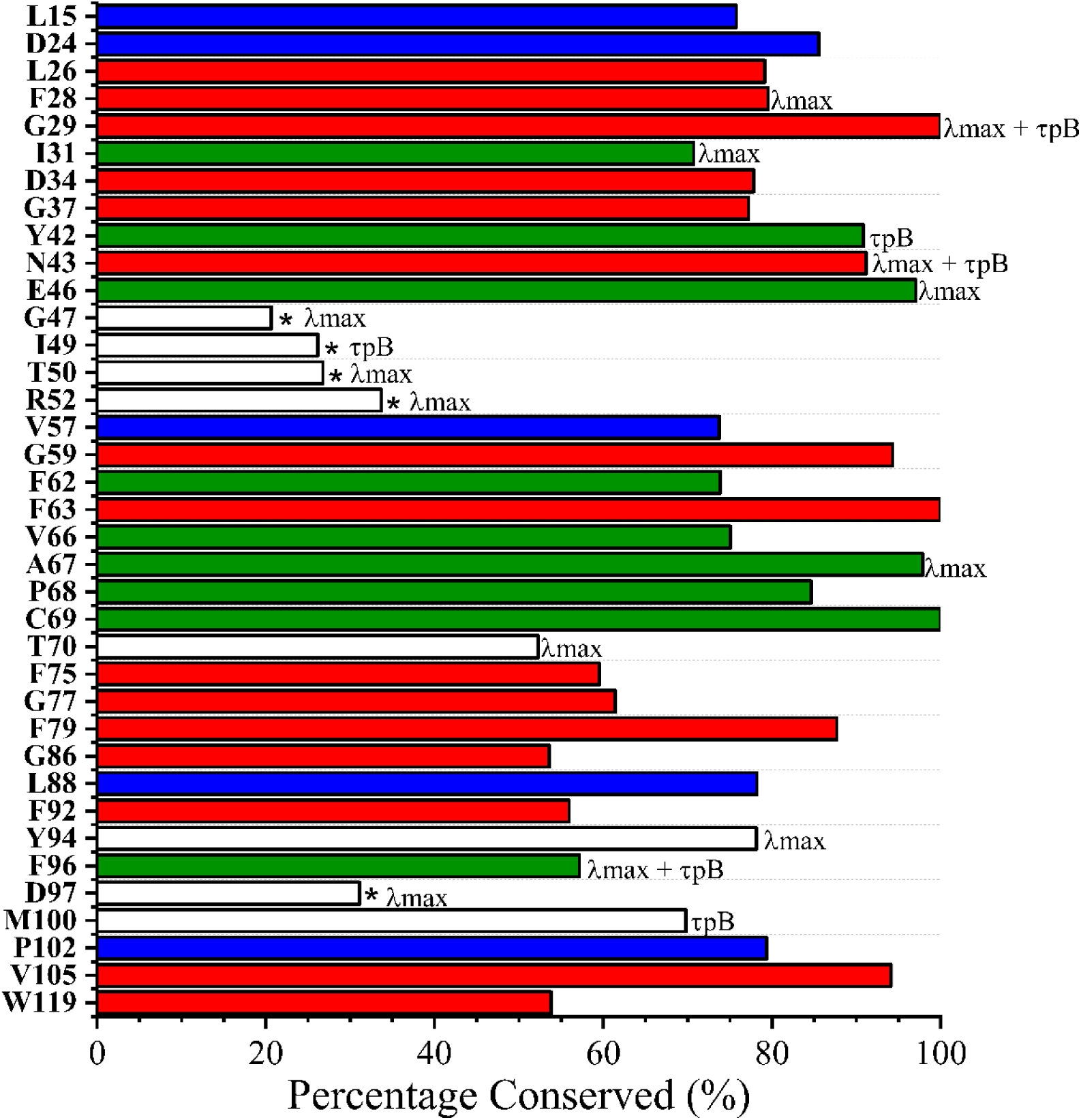
Level of conservation of important functional residues of PYP. For a set of 984 PYP sequences the level of conservation was determined using residue numbering based on Hhal PYP 1. Indicated are residues in the conserved *p*CA binding pocket (green), other conserved residues determined by Kumauchi et al,. 2008 (8) (red), residues that showed large effects in the alanine scan of Hhal PYP 1(white), and new conserved residues identified here (blue). Based on the data from the alanine scan, the residues were marked accordingly, the absorbance maximum shifted (λ_max_), the photocycle rate was affected (τ_pB_) and if both propertied were affected it is annotated as such. For the five residues marked with an * the residue present in Hhal PYP is not the consensus for the set of PYPs studied here.

First, we examined the degree of conservation of residues that were previously identified (8) as forming a conserved binding pocket (green bars in Figure 4): Ile31, Tyr42, Glu46, Phe62, Val66, Ala67, Pro68, Cys69 and Phe96. Based on studies of Hhal PYP, Tyr42, Glu46, and Cys69 would be expected to play an indispensable role. Cys69 forms the thioester bond that covalently links the *p*CA chromophore to PYP (17, 18, 20, 21). Glu46 forms an ionic hydrogen bond to the *p*CA (45) and serves as the proton donor for the *p*CA during the PYP photocycle (25), which is essential for driving conformational changes upon formation of the pB intermediate (25, 57). 97.1% of the homologs studied here contain Glu46, while the remaining 2.6% exhibit Gln46. The E46Q mutant of Hhal PYP reduces the structural changes when the pB signaling state is formed (78, 79), and the functional properties of the naturally occurring PYPs containing Gln46 remain to be studied. In Hhal PYP, the Y42F mutation causes substantial perturbations to the protein, including a thermal equilibrium between species absorbing near 446 nm and near 400 nm (76, 80–82). The hydrogen bond between Tyr42 and the *p*CA chromophore is therefore considered to be important for the stabilization of the native state of the active site of PYP (83). Tyr42 is observed in 90.8 % of the PYP homologs reported here, while in the remaining cases (8.9%) it is Phe42. For the remaining six residues in a previously proposed conserved *p*CA binding pocket: Ile31, Phe62, Val66, Ala67, Pro68, Phe96 (84), we find conservation levels ranging from 73% for Ile31 to 96% for Ala67, with the single exception being Phe69, which is conserved at a lower level (56%).

Second, we examined a set of 15 residues previously identified (84) to be >85% conserved in PYP (red bars in Figure 3): Leu26, Phe28, Gly29, Asp34, Gly37, Asn43, Gly59, Phe63, Phe75, Gly77, Phe79, Gly86, Phe92, Val105 and Trp119 (8). Using a cutoff value of >70% sequence identity, we arrive at an updated set of highly conserved residues in the PYP family (Leu26, Phe28, Gly29, Ile31, Asp34, Gly37, Tyr42, Asn43, Glu46, Gly59, Phe62, Phe63, Ala67, Pro68, Cys69, Phe79, Val105), with Phe75, Gly77, Gly86, Phe92, Phe96, and Trp119 exhibiting lower levels (54-61%) of conservation. In addition, the larger set of diverse PYPs reported here reveal Leu15, Asp 24, Val57, Leu88, and Pro102 as residues that are >70% conserved (blue bars in Figure 4).

Third, we examined the set of 16 residues that caused the largest changes (as indicated in Figure 4) in λ_max_ or τ_pB_ (or both) in the complete mutagenesis scan of Hhal PYP (67). Seven of these residues are quite strongly conserved, with 5 being in the *p*CA binding pocket. However, six of these residues (Gly47, Ile49, Thr50, Arg52, Thr70, Asp97) exhibit fairly low levels of conservation (from 21% to 53% sequence identity), and for positions 47, 49, 50, 52, and 97 (indicated with an asterisk in Figure 3) the residue present in Hhal PYP does not represent the consensus in the set of PYPs reported here. Instead, the consensus residues for the PYP family derived here are: Ser47, Leu49, Ala50, Leu52, and Thr97. A consensus sequence of PYP is depicted in Supplemental Figure S1. These six residues, together with Ile31, Phe96 and Met100 (see Figure 4), are candidates for residues that are used in nature to contribute to the wide range of values of λ_max_ or τ_pB_ observed in PYP homologs. These considerations also indicate that residues Phe28, Gly29, Tyr42, Asn43, Glu46, Ala67 and Tyr94, while causing large changes in functional properties upon mutation in Hhal PYP, are highly conserved and thus unlikely to be used in nature to tune the properties of most PYPs.

### Identification of functionally related protein as part of a multidomain protein

83% of the Cys69-containing PYP homologs reported here occur as single domain protein, showing that this is the preferred mode of *in vivo* PYP function. The remaining 17% were observed to be larger than 200 amino acids and were classified as multi-domain proteins. Interestingly, the flavin-based blue-light photoreceptor LOV, which also is a member of the PAS domain superfamily, predominantly occurs as part of multi-domain protein (85). By analyzing the PYP-containing multidomain proteins, we aimed to identify potential components of PYP signal transduction chains and to obtain insights into proteins that may be functionally related to PYP.

Using the NCBI Conserved domain database (70), we identified various domains within PYP-containing multi-domain proteins (Figure 5). Among these proteins, 22% contained a phytochrome domain followed by histidine kinase domains (HisKA, HWE-HK and HATPase_c) domains (37 cases), while 9% included a phytochrome domain followed by GGDEF/EAL domains (15 cases). In all of these proteins, PYP was present as the N-terminal domain (Figure 5A). These results build on the initial identification of a PYP-Phy-HisKA protein in *Rhodospirillum centenum* (39) and of a PYP-Phy-GGDEF/EAL protein in *Thermochromatium tepidum* (42, 43) and demonstrate that PYP-containing proteins with this domain architecture occur frequently. These multi-domain proteins are predicted to integrate both blue- and red-light photoreceptor functionalities and contain signal relay domains involved in protein phosphorylation and regulating the cytoplasmic concentration of c-d-GMP, respectively. For comparison, the LOV-containing multidomain protein PHY3 in *Adiantum capillus-veneris* includes a phytochrome domain followed by two LOV domains and a C-terminal serine threonine kinase domain, enabling phototropism and chloroplast movement in response to red and blue light (86). Such PYP-Phytochrome multi-domain proteins with signaling components like GGDEF/EAL or HisKA/HATPase may mediate various potential color-sensitive responses to light: light-regulation of gene expression or biofilm formation, negative phototaxis under potentially harmful blue light, and positive phototaxis under red light that is favorable for photosynthesis. Experimental validation is required to determine the biological response triggered by these proteins and the exact function of each component of these multi-domain proteins. Proteins with this domain architecture are primarily found in the families Methylobacteriaceae (facultative methylotrophs) and Chromatiaceae (purple sulfur bacteria and photolithoautotrophs).

**Figure 5.**
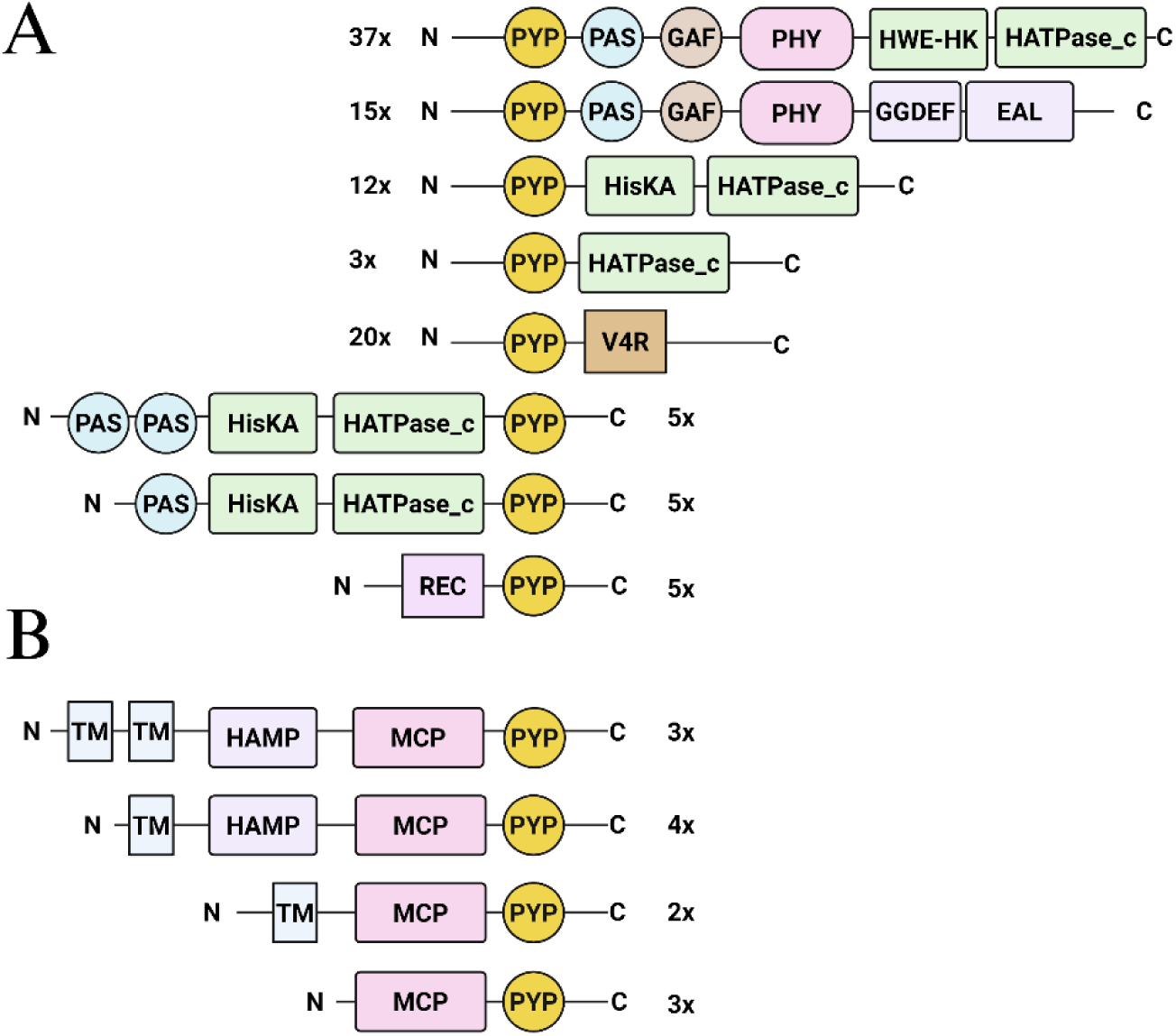
Domain composition of multi-domain proteins containing PYP. (A) PYP indicates photoactive yellow protein, PAS is a Per-Arnt-Sim domain, PHY is a phytochrome domain, HisKA/ HATPase_c is a histidine kinase domain, GGDEF/EAL perform synthesis and breakdown of c-di-GMP, V4R is a small molecule binding domain and REC is a TCRC receiver domain. Both proteins containing PYP at the N-terminal end (A) and the C-terminal end (B) were observed. (B) domain structure of MCP-PYP fusion proteins. TM indicates a predicted transmembrane α-helix.

In 15 proteins, PYP was also found to be located at the N-terminal end of shorter multi-domain proteins containing either a combination of histidine kinase and HATPase_c domains or a single HATPase domain. The HATPase domains are associated with a variety of functions, including histidine kinases, and are potentially components of a TCRS. Thus, the predicted mode of action of these proteins is that PYP serves as a blue light input domain while the HATPase domain relays the signal to an associated response regulator.

We found that PYP is also fused to a vinyl-4-reductase (V4R) domain in 20 different multi-domain proteins, where it is consistently located at the N-terminal regions. The V4R domain has been proposed to bind small molecules, potentially hydrocarbons (87), and for selected cases this prediction has been confirmed (88). These previously unidentified PYP-V4R fusion proteins thus appear to combine input domains that respond to blue light and to the presence of a small molecule ligand. We also observed five cases of a Rec-PYP fusion protein. Since Rec domains generally are the phosphorylated domain of the second component of TCRSs, this protein shares the property of the PYP-V4R proteins that it appears to combine two signal input domains while lacking an apparent signal relay domain.

While PYP is predominantly located at the N-terminal end of multi-domain proteins (see Figure 5), we observed multiple instances of proteins containing PYP at the C-terminal end, including proteins with a PAS-PAS-HisKA-HATPase_c-PYP domain structure (Figure 5A). The most notable set of proteins containing a C-terminal PYP domain are 12 cases of PYP fused to a methyl-accepting chemotaxis protein (MCP) (Figure 5B). MCP proteins have been studied extensively as the receptors that initiate chemotaxis in *E. coli* (56). The identification of 12 PYP- MCP fusion proteins provide a strong validation of the first instance of a protein with such a predicted multi-domain structure in *Methylomicrobium alcaliphilum* (11). The combination of photoreceptor and MCP functionalities is reminiscent of the signal relay mechanism for the sensory rhodopsins in *Halobacterium salinarum*, where the transmembrane SR photoreceptor forms a complex with a transmembrane MCP homolog called Htr that relays the biological signal into a ChA/CheY-like pathway that results in phototaxis (89). This resemblance indicates the possibility that the MCP-PYP multidomain protein relays phototactic signals through the CheA/CheY pathway, initiating flagellar motor rotation. Structural predictions using AlphaFold support the notion that the MCP moiety in these fusion proteins adopts a classical MCP-like conformation (Figure 6). Interestingly, we detected MCP-PYP homologs with the classical structure of two predicted transmembrane alpha-helices and a HAMP domain, but also MCP- PYPs with zero or one predicted transmembrane alpha-helices or that lacked the HAMP domain (90). Functional MCP homologs lacking transmembrane helices were reported in *Sinorhizobium meliloti* (91).

**Figure 6.**
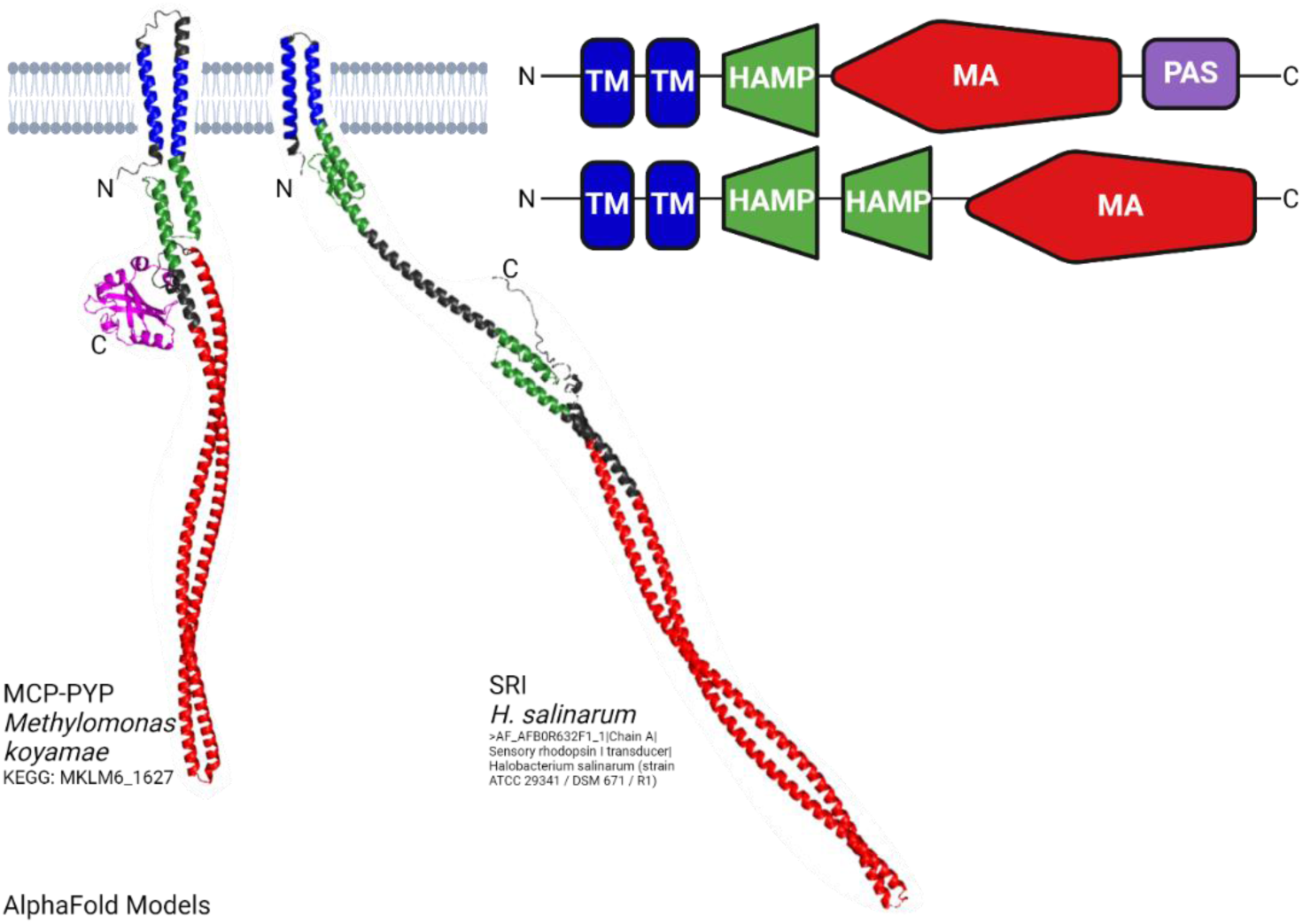
Predicted structures of the MCP-PYP multi-domain protein from *Methylomonas koyamae* and HtrI from *Halobacterium salinarum*. Left: predicted three-dimensional structures of these two MCP homologs. Right: derived domain architectures, with the color coding of the domains allowing direct comparison with the predicted three-dimensional structures. The PYP domain is indicated in purple. HtrI contains an additional HAMP domain.

The set of 153 multi-domain proteins reported here shows that PYP is preferentially located at the N-terminal region (65% of cases), while in 35% of cases it is found at the C-terminal end (Figure 5). Interestingly, both LOV shows the same preference for N-terminal localization (85). The placement of the PYP domain at the N- or C-terminal end varied strongly depending on the domain composition of the protein. For all 52 PYP-PHY multi-domain proteins and all 20 PYP-V4R proteins, the PYP domain was located at the N-terminus. In contrast, PYP showed a C-terminal location for all 12 MCP-PYP fusion proteins (Figure 5). In the case of HisKA/HATPase domains, the location of PYP was mixed.

One possible explanation for these observations is that the preferential N- or C-terminal position of the PYP domain is dictated by the molecular mechanism of inter-domain signal relay within the multi-domain protein. While intriguing, this possibility will require further research. First, it should be pointed out that it is difficult to fully exclude the possibility that the observed domain architecture is not caused by such a selective pressure, but because of inheritance, in which a specific multi-domain architecture arose once and then was inherited by horizontal and/or vertical gene transfer. We examined if the similarity of PYPs, their domain structure, and the taxonomy of the organism in which the PYP is present are related and detected clear patterns in this complex relationship of biological properties (Figure 7). However, no straightforward predictions on the function of PYPs were obtained from this analysis.

**Figure 7.**
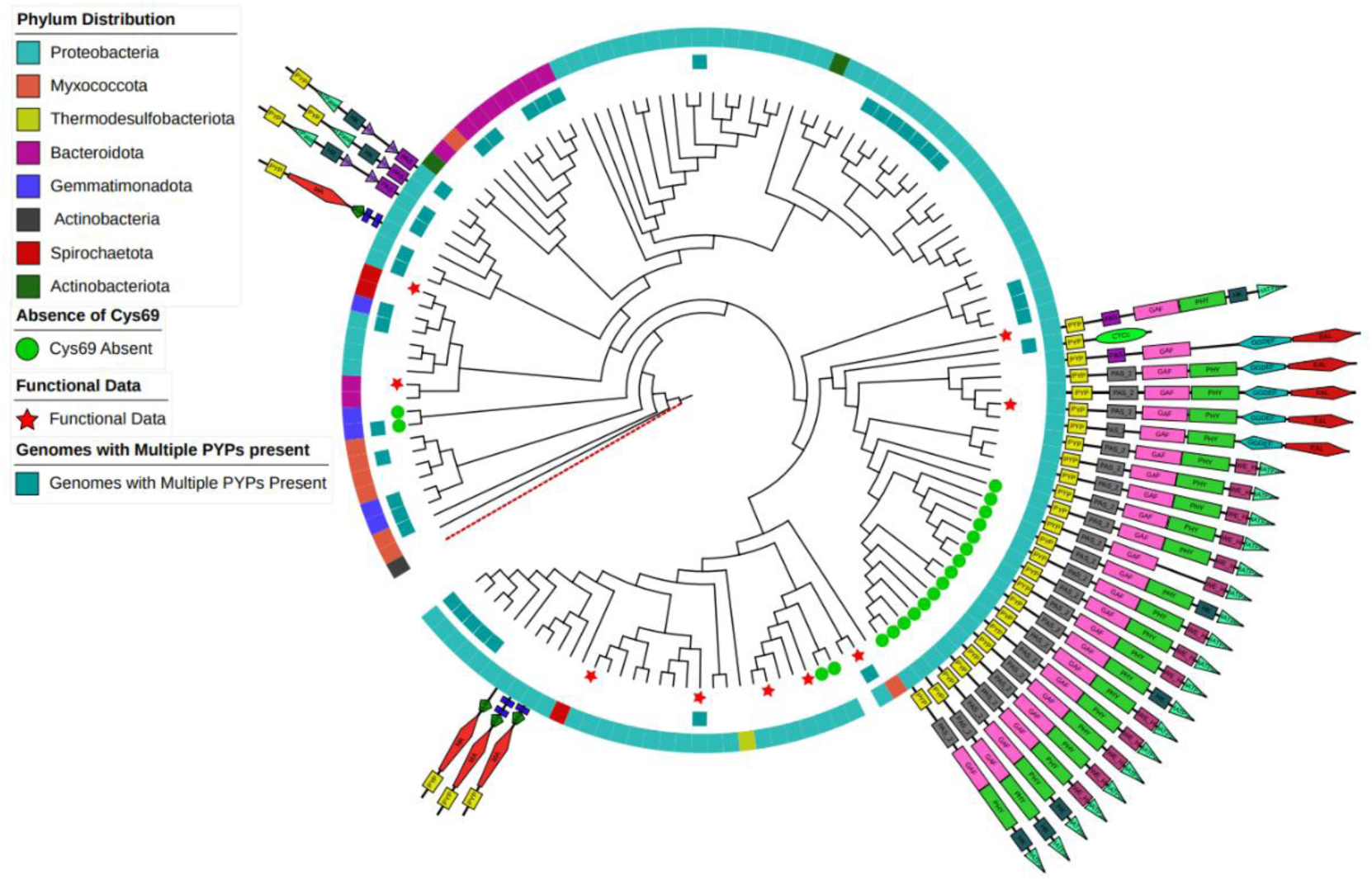
Depiction of selected properties of the PYPs reported here. The phylogenetic gene tree depicts the relationship of the *pyp* homologs. This tree was computed based on only PYP, also if a PYP domain was analyzed.

Second, in a question of more general relevance for the field of biological signal transduction, it remains to be determined how the location of a domain in a multi-domain protein affects signaling. Many PYPs (and some LOVs) are proficient in signaling as single-domain proteins, demonstrating that signal relay from these proteins can proceed even if they are not part of a multi-domain protein. In those cases, signal relay presumably occurs via transient or stable (as in the case of SR-Htr protein complexes) protein-protein complexes. The advantage of incorporating PYP into a multi-domain protein remains to be determined.

### Experimental studies related to the PYP-MCP multi-domain protein from Nitrincola alkalilacustris

*Nitrincola alkalilacustris* DSM-No Strain 29817 is one of the 12 organisms identified here containing a PYP-MCP multidomain protein. This protein provides an attractive candidate for examining the molecular signal transduction mechanism of phototaxis predicted to be initiated by this PYP homolog. We first examined if this organism exhibits motility and used electron microscopy to examine if *N. alkalilacustris* cells have flagella (see supplement for a description of the experimental methods). The results confirmed the presence of flagella on these bacteria (Figure 8A). We also detected motility on low percentage agar plates (Figure 8B), confirming the results from the TEM images. To examine if the PYP domain in the MCP-PYP protein binds *p*CA, is yellow and exhibits photochemical activity, we overexpressed it in *E. coli*. The resulting purified protein was studied using UV/Vis absorbance spectroscopy, demonstrating that the PYP from *N. alkalilacustris* (Nalk PYP) has an absorbance maximum at 447nm (Figure 8D), identical to the well-studied PYP from Hhal. The extinction coefficient of Nalk PYP is 43.8 mM^-1^ cm^-1^, very similar to the value of 45.5 reported for Hhal PYP (92). We used time-resolved UV/Vis absorbance spectroscopy to examine if Nalk PYP exhibits photochemical activity.

**Figure 8.**
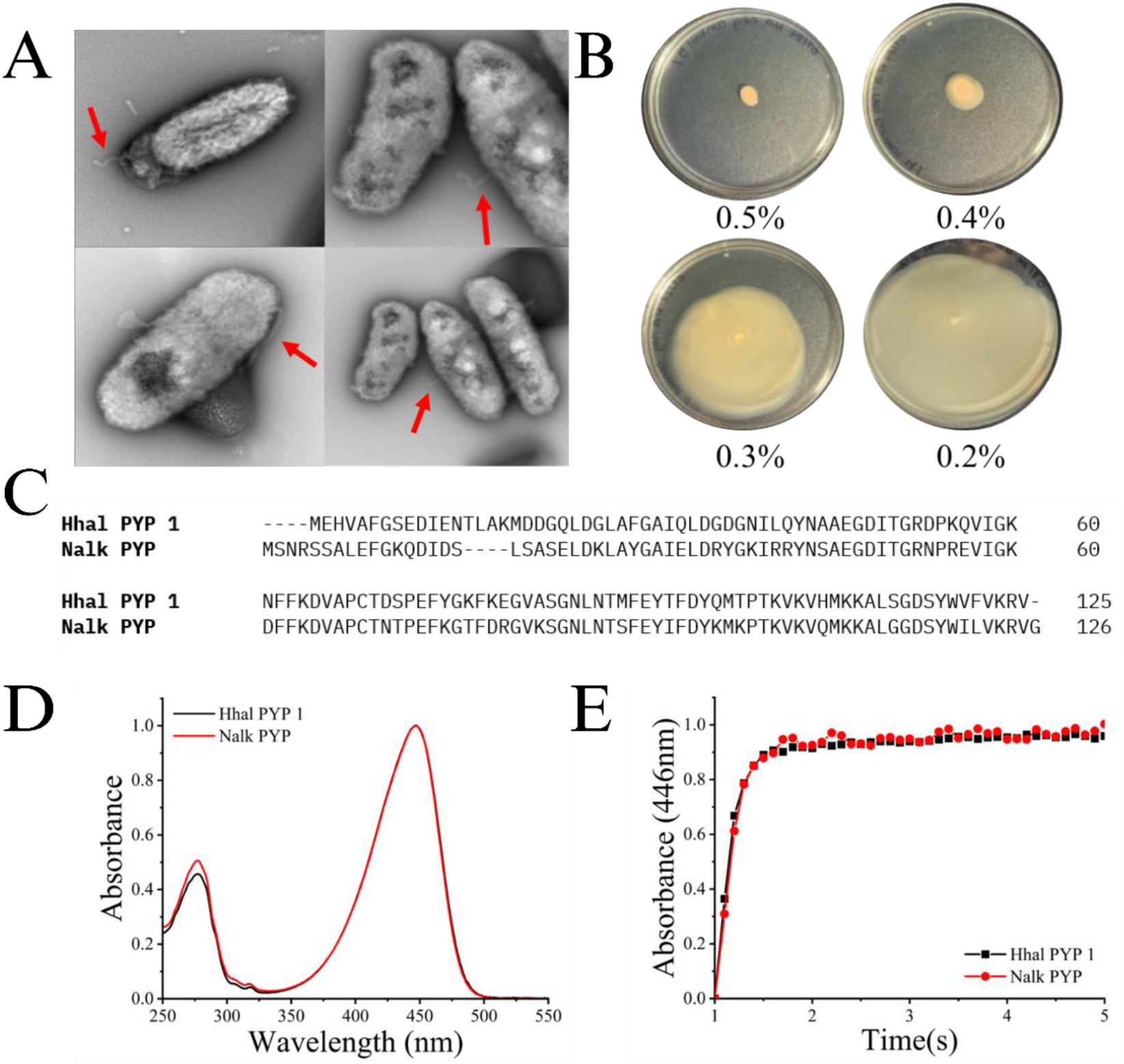
Motility of *Nitrincola alkalilacustris* DSM-No Strain 29817 and analysis of the PYP domain in the MCP-PYP multi-domain protein encoded in its genome. (A) TEM images showing flagella (indicated by the red arrows). (B) *N. alkalilacustris* grown on various agar concentrations (indicated below each plate) to examine motility in this organism. (C) Sequence alignment of Hhal PYP 1 and Nalk PYP. (D) UV/Vis absorbance spectrum of *N. alkalilacustris* PYP in comparison to the well-studied PYP from *Halorhodospira halophila.* (E) Kinetics of decay of the pB photocycle intermediate after illumination in Nalk PYP (red circles) and Hhal PYP (black circles) detected by changes in absorbance at 446 nm. The data were fitted as a mono-exponential decay (solid lines).

Indeed, illumination of the pure protein resulted in the formation of a blue-shifted intermediate with an absorbance maximum near 350 nm. Decay of this pB photocycle intermediate occurred with a time constant of 0.5 seconds, essentially identical to the rate of pB decay for Hhal PYP (Figure 8E). Thus, both the λ_max_ and τ_pB_ of Nalk PYP are identical to those of Hhal even though their sequences vary considerably. Nalk PYP shares 60.8% sequence identity with Hhal PYP (Figure 8C). The lifetime of the pB intermediate of Nalk PYP is consistent with a role of this protein in triggering phototaxis. In line with this possibility, the genome of *N. alkalilacustris* encodes homologs of all components needed to form a functional chemotaxis signal transduction pathway (Supplemental Table S1). However, thus far we have not able to detect phototaxis responses in this organism.

### Analysis of predicted pyp operons

We aimed to complement the search for candidate proteins that are functionally related to PYP based on the above analysis of multi-domain PYPs with an examination of genes present in predicted operons containing the *pyp* gene. Below, we will refer to these predicted operons as *pyp* operons. The first *pyp* operon was identified in *Halorhodospira halophila* (37, 52, 53). This operon consists of three genes: the *pyp* gene followed by a gene with clear homology to *p*- coumaryl CoA ligase (*p*CL) and a gene with clear homology to tyrosine ammonia lyase (TAL). The proteins encoded by these genes were experimentally demonstrated to indeed have pCL and TAL activity (53, 93, 94). The activity of TAL is to convert tyrosine to *p*CA, and that of *p*CL is to use ATP to activate *p*CA by attaching it to Coenzyme A. The resulting *p*-coumaryl-CoA then reacts with apoPYP to yield functionally active holoPYP in which *p*CA is covalently attached to Cys69.

Given that *p*CA is essential for PYP function, we expected that the majority of *pyp* operons would contain genes encoding TAL and *p*CL. However, our analysis of 113 different *pyp* operons demonstrated that this is not the case. A gene encoding TAL was found in only 5% of the predicted *pyp* operons (7 instances), while 32 % of *pyp* operons (42 instances) contained a gene encoding *p*CL. These results suggest that genes encoding enzymes with TAL activity are located elsewhere in the genome of these organisms. It is also possible that some of the bacteria involved lack TAL activity and are able to take up *p*CA from their environment. We examined the degree of sequence conservation in the 7 TAL enzymes encoded in the *pyp* operons analyzed here and found that they exhibit strong sequence divergence (Supplemental Table S2), complicating their conclusive functional annotation (particularly with respect to substrate specificity). Similarly, genes encoding the *p*CL activity necessary to yield holoPYP are likely to be encoded elsewhere in the genome of these organisms. It has been reported that genes encoding predicted TAL and *p*CL enzymes can be located in the vicinity of *pyp* gene (11). The enzymes involved can have a broad substrate range (95–98) and may be annotated as ammonia lyases or CoA ligases for substrates other than tyrosine or *p*CA, rendering bioinformatics analyses of the presence of these genes more challenging.

A distinct set of recurring proteins was found to be encoded in genes that are part of *pyp* operons (Table 1): GGDEF/EAL proteins (in 29 *pyp* operons) and HisKA and HATPase (in 17 operons). We also observed two instances in which a *pyp* operon encoded a BLUF protein. Since BLUF is a flavin-based blue-light photoreceptor (99, 100), these predicted operons encode two different types of blue-light photoreceptors. It should be noted that both BLUFs were found in *pyp* operons in the genome of the same organism (Caballeronia sp. SBC2). This genome also encodes 5 different *pyp* homologs.

**Table 1.**
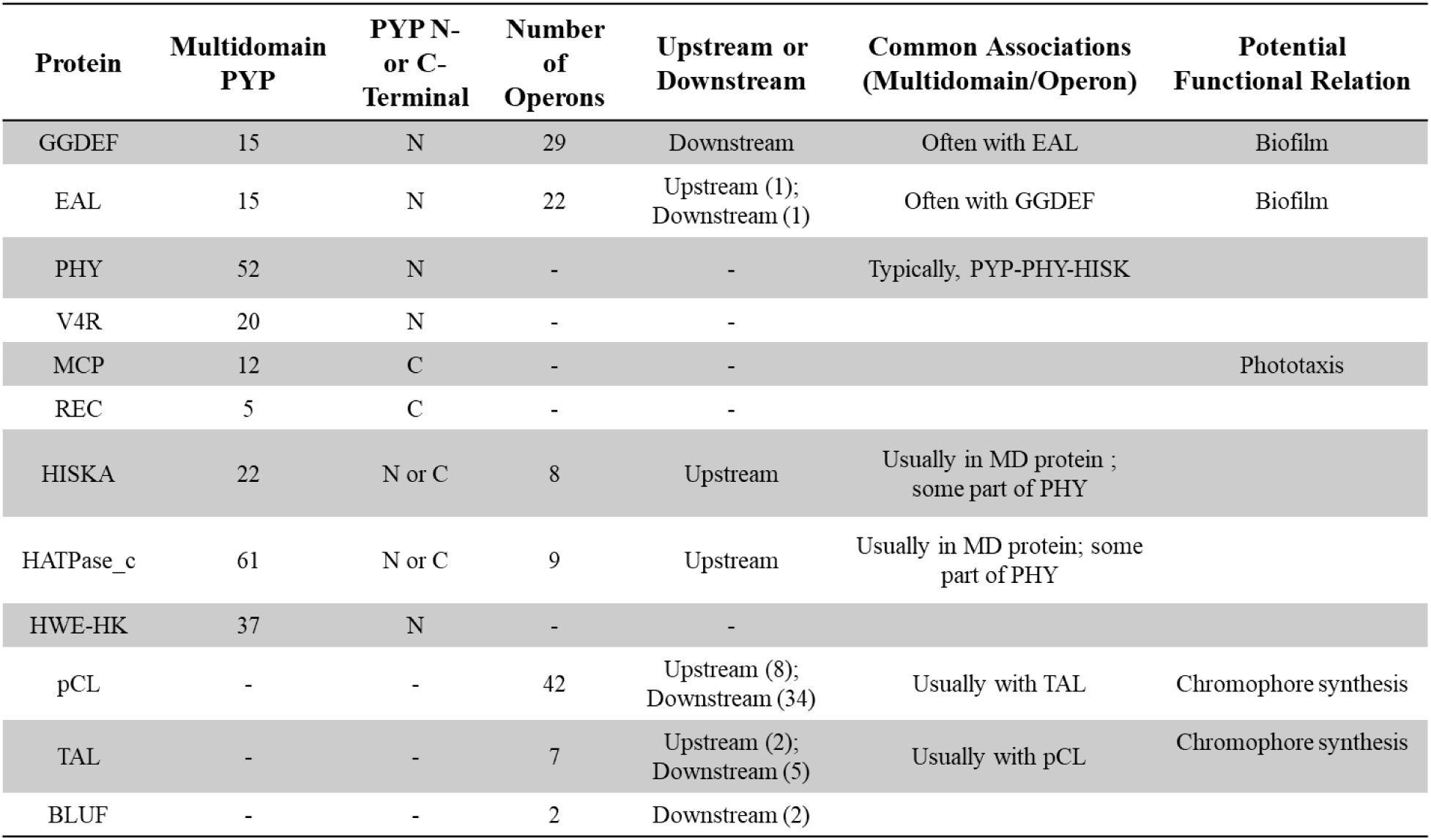
A list of commonly occurring proteins found both in an operon and/or in a multidomain protein with PYP. The proteins are annotated for where they are in regard to PYP.

| Protein | Multidomain PYP | PYP N- or C-Terminal | Number of Operons | Upstream or Downstream | Common Associations (Multidomain/Operon) | Potential Functional Relation |
| --- | --- | --- | --- | --- | --- | --- |
| GGDEF | 15 | N | 29 | Downstream | Often with EAL | Biofilm |
| EAL | 15 | N | 22 | Upstream (1); Downstream (1) | Often with GGDEF | Biofilm |
| PHY | 52 | N | - | - | Typically, PYP-PHY-HISK |  |
| V4R | 20 | N | - | - |  |  |
| MCP | 12 | C | - | - |  | Phototaxis |
| REC | 5 | C | - | - |  |  |
| HISKA | 22 | N or C | 8 | Upstream | Usually in MD protein ; some part of PHY |  |
| HATPase_c | 61 | N or C | 9 | Upstream | Usually in MD protein; some part of PHY |  |
| HWE-HK | 37 | N | - | - |  |  |
| pCL | - | - | 42 | Upstream (8); Downstream (34) | Usually with TAL | Chromophore synthesis |
| TAL | - | - | 7 | Upstream (2); Downstream (5) | Usually with pCL | Chromophore synthesis |
| BLUF | - | - | 2 | Downstream (2) |  |  |

We observed a striking preference for the genetic association of some proteins either based on their presence in a PYP-containing multi-domain protein or encoded in a *pyp* operon (Table 1). While PYP-Phy fusion proteins were the most abundant type of PYP-containing multi-domain proteins, we did not observe a single instance of a *pyp* operon encoding a phytochrome homolog. In the case of both GGDEF/EAL and HisKA/HATPase proteins, both genetic association with PYP in multi-domain protein and in the same operon was observed. For the genes encoding HisKA/HATPase proteins, we noticed a preference for being present upstream of the *pyp* gene. At this point, insufficient information is available to provide mechanistic explanations for these observations.

### Organisms with genomes encoding with multiple PYP homologs

*H. halophila* is the organism in which the first PYP was identified (9, 10), and is the first – and currently only – organism reported to encode more than one PYP (44). Notably, 16% of the genomes analyzed here encoded more than one PYP homolog (supplemental Table S3). This observation shows that the case of two *pyp* homologs encoded in one genome is not unique to *H. halophila*. The phototaxis of responses *Halobacterium salinarum* are governed by sensory rhodopsin I and II, providing a classic example of two homologous photoreceptors that trigger distinct positive and negative phototaxis responses towards different colors of light (89). However, since the photocycle rate of Hhal PYP 2 is much slower than that of Hhal PYP 1 (approximately 50 seconds versus approximately 0.5 seconds), it is also possible that some of the PYP homologs in these organisms trigger biological responses other than phototaxis, but this question awaits further experimental work.

### Insights into proteins functionally associated with PYP and its biological functions

Experimental evidence has been reported that PYP can trigger three distinct biological functions: negative phototaxis in *H. halophila* triggered by a single-domain PYP via an unknown signal transduction chain (33), light-regulation of enzymes that produce photoprotective pigment biosynthesis by a PYP-Phy-HisK multi-domain protein, where light-regulation of the HisK domain causes signaling via a TCRS (39), and blue-light regulation of biofilm formation via a GGDEF-based signaling mechanism (48).

The results reported here provide clear support for all three of these signaling mechanisms. First, the occurrence of MCP-PYP fusion proteins in multiple bacteria strengthens the occurrence of PYP as a blue light receptor for phototaxis and indicates that the signal transduction chain linking PYP to the flagellar motor involves components of the classical *E. coli* chemotaxis signaling machinery. The lifetime of the pB intermediate of Nalk PYP of 0.5 seconds is consistent with this proposal. These results also suggest the possibility that the PYP from *H. halophila* triggers negative phototaxis by interacting with an MCP or other component of the chemotaxis machinery. Second, the genetic link between PYP and HisKA/HATPase, observed both in PYP-containing multi-domain proteins and in *pyp* operons, indicates that PYP can relay signals into an associated TCRS, as was reported for the PYP homolog from *Rhodospirillum centenum* (39). Since the biological responses triggered by a TCRS can vary greatly and are difficult to predict based on sequence information, this result does not provide a specific hypothesis about the biological function of these PYPs. Third, the genetic link between PYP and GGDEF/EAL proteins observed both in PYP-containing multi-domain proteins and in *pyp* operons indicates that PYP can be involved in the blue-light regulation of biofilm formation by modulating cellular c-di-GMP content (101). These results provide testable hypotheses regarding the biological function and signaling mechanisms of PYP homologs that can be experimentally verified. In addition, the presence of PYP in proteins that also contain a phytochrome domain, the finding of PYP-V4R proteins reported here, and the observation that multiple organisms contain more than one *pyp* gene indicate that many aspects of the bacterial photobiology associated with PYP, including the integration of multiple light and chemical signals, remain to be explored.

## Acknowledgements

The authors thank Dr. Chelsea Murphy at the High-Performance Computing Center at Oklahoma State University for help with various bioinformatics analysis, the support team at the Microscopy core of facility Oklahoma State University, and Dr. Matthew Cabeen for helpful suggestions on the text of this manuscript.

